# Computational assay of hERG ion permeation and its modulation by small-molecule compounds

**DOI:** 10.64898/2026.10.21.760785

**Authors:** Ivan Manoza, Vytautas Gapsys, Wojciech Kopec

## Abstract

The human ether-a-go-go-related gene (hERG) potassium channel is an important off-target in drug discovery, as its inhibition can disrupt cardiac repolarisation and cause potentially fatal arrhythmias. Despite its importance, the molecular mechanisms underlying hERG ion permeation and its modulation by small molecules remain elusive. Here, we use molecular dynamics simulations to reproduce sustained K^+^ permeation and single-channel conductance comparable to experimental measurements. Our simulations reproduce the characteristic asymmetry in hERG single-channel conductance and reveal a structural mechanism underlying this behaviour. We identify the conformational dynamics of S624, located at the innermost ion-binding site of the selectivity filter, as a determinant of this conductance asymmetry. Substitution of S624 with the canonical threonine abolishes conductance asymmetry by facilitating K^+^ entry into the S4 binding site. Simulations of classical hERG inhibitors from cryo-EM-derived binding poses show that bound inhibitors sterically occlude K^+^ access to the selectivity filter, with binding stabilised by interactions with hydrophobic residues lining the central cavity. In contrast, simulations of hERG activators support a fenestration binding mode consistent with mutagenesis data, in which PD-118057 and PD-307243 increase cavity K^+^ occupancy and enhance conductance by two- to threefold relative to the apo channel. Together, these results establish a molecular framework connecting hERG structure, ion permeation, and small-molecule modulation, providing a basis for directly assessing the functional consequences of compound binding.

## 2. Introduction

Potassium channels are present in most cell types, where they regulate a range of essential cellular functions and play an important role in restoring the membrane potential following depolarisation. All K^+^ channels contain a selectivity filter (SF) composed of four adjacent ion-binding sites, primarily formed by backbone carbonyl oxygens (S1– S3), with the S4 site typically involving hydroxyl oxygens from threonine residues.^1^ The human ether-a-go-go-related gene (hERG) channel, a member of the voltage-gated potassium (K_V_) family,^2^ is particularly important in cardiac cells, where it controls ventricular repolarisation and determines the QT interval on the electrocardiogram.^3^

While hERG shares a K^+^-selective pore and four voltage sensors with other K_V_ channels, it exhibits several distinctive features. First, its SF contains an atypical amino acid sequence, Ser-Val-Gly-Phe-Gly-Asn, in place of the canonical Thr-Val-Gly-Tyr-Gly-Asp found in most K^+^ channels.^1^ Second, it displays low single-channel conductance relative to other K_V_ channels.^4^ Third, hERG exhibits inward rectification; however, unlike classical inward rectifier channels, where rectification arises from intracellular block by divalent ions and polyamines,^5,6^ hERG rectification has been linked to rapid C-type-like inactivation of the SF.^7,8^ However, this asymmetry also exists at the level of single-channel conductance, with inward currents consistently reported to be two- to fourfold larger than outward currents.^9^

The clinical significance of hERG became clear when drugs were withdrawn from the market after causing fatal cardiac arrhythmias through unintended hERG channel block,^10^ leading to prolongation of the QT interval, a condition known as drug-induced long QT syndrome (di-LQTS).^11,12^ As a result, screening potential drug candidates for hERG inhibition has become a key requirement in drug development.^13^ Early *in silico* work by Cavalli *et al.* identified a shared pharmacophore among hERG blockers, characterised by a protonated basic amine and one or more aromatic moieties.^14^ Mutagenesis studies identified residues critical for blocker binding, implicating F656, Y652 and G648 on the S6 helix, and T623, S624 and V625 on the P-loop as key determinants.^15^ These residues line the central cavity, marking it as the primary binding site for hERG blockers.^16^ More recently, higher-resolution cryo-EM structures of hERG bound to the classical inhibitors astemizole, E-4031 and pimozide revealed that all three inhibitors adopt a horizontal orientation beneath the SF.^17^

In contrast to blockers, a small number of hERG activators have been shown to increase K^+^ conductance through hERG. These compounds increase outward K^+^ current by reducing the extent of inactivation, altering its voltage dependence, or prolonging the open state through slower deactivation.^18^ A set of activators known as negatively charged activators (NCAs) shares a common pharmacophore comprising a negatively charged aromatic moiety and a hydrophobic moiety.^19^ The binding mode of NCAs within the channel remains a matter of debate. One proposed site is beneath the SF, where their negatively charged group coordinates a K^+^ ion, increasing SF occupancy and promoting channel opening.^19^ An alternative binding mode positions the activators deeper within the hydrophobic fenestration site at the side of the central cavity, which better accounts for the mutagenesis data.^20^ Nevertheless, sustaining ion permeation in hERG over long timescales remains a long-standing challenge, making validation of competing binding modes and their functional effects through molecular dynamics (MD) particularly difficult.

Molecular dynamics simulations under applied voltage (sometimes referred to as *in silico* or computational electrophysiology) provide an unfiltered view of ion permeation in ion channels, including the direct knock-on mechanism in potassium channels^21^ and the molecular basis of ion selectivity.^22^ However, previous simulations of hERG showed a highly unstable SF, which rapidly deformed from its conductive configuration and prevented sustained ion permeation.^23^ Permeation events were observed only at an unrealistically high applied voltage of 750 mV, yielding a conductance estimate of *∼*0.8 pS, well below the experimental value of 3–5 pS.^4,9,23^ While fast inactivation is a characteristic feature of hERG, the SF should remain sufficiently stable over hundreds of nanoseconds to sustain ion permeation; otherwise, no macroscopic current would be observed.

More recently, force fields based on the electronic continuum correction (ECC) approach have led to substantial improvements in simulating ion permeation and SF ion occupancy in K^+^ channels.^24^ ECC-based simulations of three K^+^ channels brought simulated single-channel conductance, SF ion occupancy and K^+^/Na^+^ selectivity into close agreement with experimental measurements.^24^ Notably, however, all channels tested to date share a canonical SF sequence, and whether ECC can similarly benefit potassium channels with a non-canonical SF such as hERG has not yet been investigated.

In this work, we address the long-standing challenge of simulating ion permeation in hERG by combining two approaches: the stabilising, non-inactivating N629D mutation at the outer mouth of the SF and recently developed ECC-based force fields. Together, these approaches enable, for the first time, robust microsecond-timescale CompEL simulations of hERG-N629D under applied voltage (*±*300 mV), in which the SF remains stable and supports sustained ion permeation. Our findings reproduce key experimental properties of hERG, including single-channel conductance in qualitative agreement with experiment and its characteristic voltage-dependent asymmetry, with the S4 site of the SF emerging as a possible structural basis for the latter.

Building on this, we investigate the modulation of ion permeation by small molecules. Simulations of the hERG inhibitors astemizole, E-4031 and pimozide provide mechanistic insight into how these compounds block hERG conduction. Finally, we test the proposed binding modes of the NCAs PD-118057 and PD-307243. Our findings demonstrate that fenestration-bound activators increase ion permeation and remain stable, maintaining key contacts with L646 and F619. This study examines sustained hERG ion permeation and its modulation by small molecules using atomistic simulations.

## 3. Results and Discussion

### 3.1 Mechanistic Basis of Asymmetric Conductance in hERG from CompEL Simulations

We performed MD simulations of hERG-N629D starting from the cryo-EM structure (PDB: 9CHP) using the ECC-Amber14SB force field, with applied voltages of *±*300 mV and ten independent replicas (1 *µ*s each) per voltage (**Figure 1a, b**). Our attempts to simulate WT hERG under the same conditions resulted in no K^+^ conduction, consistent with a recent report showing that WT hERG remains non-conductive in MD simulations, including with ECC-based force fields.^25^ SF instability in WT hERG simulations has been attributed to reorientation of F627 and remodelling of the SF towards non-conductive conformations.^23,25,26^ The N629D mutation has been shown to stabilise the conductive SF conformation, closely resembling the canonical conductive structure of KcsA.^27^

**Figure 1.**
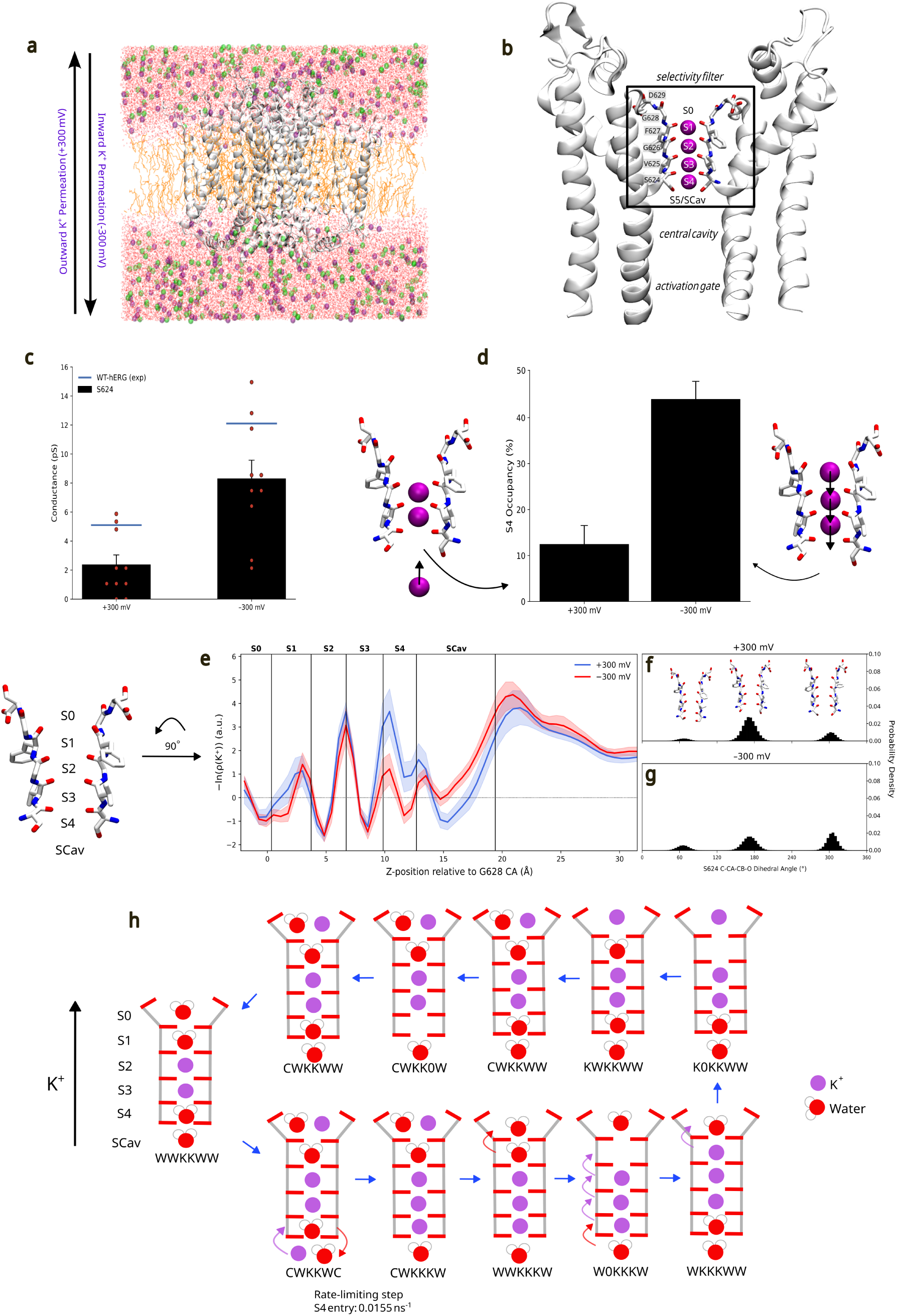
hERG-N629D CompEL simulations and mechanism of asymmetric conductance. (a) All-atom simulation system. The channel is shown in white, lipids (POPC) in orange, water as red lines, K^+^ ions as purple spheres, and Cl*^−^* ions as green spheres. A transmembrane voltage of *±*300 mV was applied, with outward and inward permeation indicated by the arrow. (b) Open-state hERG structure (PDB: 9CHP; two diagonally opposite subunits shown), highlighting the selectivity filter ion-binding sites S0–S4 and SCav. The central cavity is located below the selectivity filter and above the activation gate. Key selectivity filter-lining residues are labelled. (c) Single-channel conductance of hERG-N629D at +300 mV and *−*300 mV, with individual replica values overlaid as red dots. The blue line indicates the experimental WT hERG single-channel conductance. (d) S4 ion occupancy at +300 mV and *−*300 mV. Left inset shows a single K^+^ ion approaching S4, reflecting the absence of the multiion permeation mechanism at +300 mV. The right inset shows multiple K^+^ ions occupying the selectivity filter simultaneously, consistent with the multi-ion permeation mechanism that facilitates efficient S4 entry during inward permeation. Error bars represent SEM. (e) Potential of mean force (PMF) profiles of K^+^ along the pore axis at +300 mV (blue) and *−*300 mV (red), with shaded regions indicating standard error. Selectivity filter binding site positions are indicated by vertical lines. (f,g) Probability density distributions of the C-C*α*-C*β*-O dihedral angle of S624 at +300 mV (f) and *−*300 mV (g), with representative rotameric conformations shown above each distribution. (h) Dominant permeation cycle of apo hERG-N629D at +300 mV resolved by Kperm analysis. Each state represents the occupancy of the SF binding sites S0–S4 and SCav, where K denotes K^+^, W denotes water, C denotes co-occupancy (K^+^ and W), and 0 denotes an empty site. The rate-limiting step is ion entry into S4 from SCav, with a rate of 0.0155 ns*^−^*^1^. Error bars represent SEM.

Robust K^+^ permeation was observed at both voltages. K^+^ permeation occurred via a direct knock-on mechanism, with no co-permeating water molecules. Calculated conductance values of 8.28*±*1.3 pS and 2.35*±*0.70 pS at *−*300 mV and +300 mV, respectively, are in good agreement with experimentally reported wild-type (WT) single-channel conductance values of 12.1 *±* 0.6 pS and 5.1 *±* 0.6 pS (**Figure 1c**).^9^ Interestingly, our simulations reproduce the conductance asymmetry observed experimentally for hERG, with higher conductance at negative than at positive voltage.^9^ A similar voltage-dependent asymmetry in K^+^ permeation has recently been observed in MD simulations of hERG-N629D.^25^ In addition, this asymmetry has been observed experimentally in inactivation-attenuated mutants such as S631A,^9^ indicating a permeation-level contribution to rectification, whose structural basis remains poorly understood.

To investigate the mechanistic basis of this asymmetric conductance, we focused on the S4 binding site, which is lined by serine residues, in contrast to the majority of other potassium channels that have threonines in this position. We observed substantial conformational flexibility of the S624 hydroxyl side chain and investigated whether this behaviour influences ion permeation. Interestingly, S4 ion occupancy differs markedly between voltages: during outward permeation (+300 mV), K^+^ occupies S4 only *∼*10% of the time, whereas during inward permeation (*−*300 mV), occupancy increases to nearly 50% (**Figure 1d**). To understand the reason for this difference, we constructed an approximate free energy profile from the K^+^ density of permeating ions. A pronounced barrier is observed around the S4/SCav region (*∼*10–20 Å relative to G628 C*α*) at +300 mV (**Figure 1e**, blue trace). Analysis of the S624 C-C*α*-C*β*-O dihedral angle, which reports on the orientation of the hydroxyl oxygen, reveals a dominant population centred around *∼*180*^◦^*, where the hydroxyl oxygen points away from the pore and is unable to coordinate a permeating K^+^ ion (**Figure 1f**). We therefore propose that the dynamics of S624 contribute to the elevated free energy barrier at S4 and underlie the asymmetric conductance of the channel, i.e., lower conductance at positive voltage.

To further investigate this mechanism, we analysed ion permeation cycles using a recently published tool,^28^ and identified the dominant ion permeation pathway at +300 mV. The transition from CWKKWC to CWKKKW is the rate-limiting step, occurring with a rate constant of 0.0155 ns*^−^*^1^ (**Figure 1h**). This transition corresponds to K^+^ entry into the S4 site, which is frequently occupied by a water molecule that hydrogen bonds to a serine side chain hydroxyl group. Outward permeation, therefore, requires displacement of this water and reorientation of the S624 hydroxyl group to coordinate the incoming ion, making S4 entry the slowest step in the permeation cycle, in line with the free energy profile.

In contrast, the free energy landscape at *−*300 mV reveals a lower barrier between the S3 and S4 binding sites (**Figure 1e**, red trace). To investigate the molecular basis of this reduced barrier, we analysed the S624 C-C*α*-C*β*-O dihedral distributions. Relative to +300 mV, the rotamer population shifts towards the canonical conductive conformation (*∼*300*^◦^*), in which the hydroxyl oxygen is oriented towards the filter axis (**Figure 1f**): this corresponds to *∼*0.74 kcal/mol rotamer stabilisation in the conductive state at *−*300 mV. We therefore proposed that the conductance asymmetry arises from stabilisation of the conductive S624 rotamer, lowering the energetic barrier at S4 and increasing the probability of successful ion coordination, thereby promoting inward permeation.

This shift indicates the multi-ion permeation mechanism through the SF. During inward conduction, a K^+^ ion is ‘knocked’ into the S4 site by the strong electrostatic repulsion from ions occupying S2 and S1 sites (**Figure 1d**). While in S4, this K^+^ increases the likelihood of productive coordination with S624 hydroxyl groups. In contrast, during outward conduction, entry into S4 is essentially a single-ion process and does not benefit from the multi-ion repulsion, leaving S4 entry more dependent on the conformational dynamics of S624, ultimately resulting in a higher effective barrier for ion permeation. This distinction is reminiscent of induced fit versus conformational selection, whereby during inward permeation the ion drives productive coordination with S624, whereas during outward permeation ion entry is contingent on S624 adopting a permissive rotameric state.

To further probe the role of the S4 side chain in hERG ion permeation, we performed simulations of hERG with the S624T mutant, the canonical residue found in most K^+^ channels.^29^ Strikingly, this mutation abolished the asymmetric conductance, with S624T exhibiting similar single-channel conductance in both directions (**Figure S1a**), directly supporting the idea that S624 underlies this asymmetry. Mechanistically, we observed that T624 hydroxyl groups are restricted to only two rotameric states, strongly biasing the S4 site towards the conductive geometry (**Figure S1d,e**). We therefore examined how this reduced rotameric flexibility alters ion permeation.

At positive voltage, S4 occupancy in S624T is approximately two-fold higher relative to WT S4 (**Figure S1b**), consistent with a more favourable ion-binding geometry at this site. This is accompanied by an increase in single-channel conductance (**Figure S1a**) and a reduced free energy barrier to S4 entry (**Figure S1c**), supporting the idea that limiting S4 side chain flipping facilitates outward permeation. Permeation cycle analysis further confirms this mechanism. In WT S4, the rate-limiting step is ion entry into S4 from SCav, whereas in S624T this transition is *∼*11-fold faster and no longer a rate-limiting step. Instead, the rate-limiting step in S624T is the recruitment of K^+^ from bulk solution into SCav (0.07022 ns*^−^*^1^; **Figure S1f**). Together, these results demonstrate how restricting S4 hydroxyl groups to the conductive rotamer improves outward conductance.

At negative voltage, however, S624T exhibits reduced single-channel conductance relative to WT S4 (**Figure S1a**), despite having higher S4 occupancy than WT S4 (**Figure S1b**). The free energy profile reveals a deeper S4 minimum in S624T than in WT S4 at *−*300 mV (*∼ −*1.00 vs *−*0.76 *k*_B_*T*, **Figure S1c**), indicating that the threonine stabilises K^+^ more effectively. This stronger coordination slows exit from S4, thereby reducing net inward permeation despite higher S4 occupancy.

Collectively, these findings support a model in which the conformational flexibility of S624 limits efficient ion binding at S4 and creates a kinetic barrier to outward permeation. During inward conduction, this barrier is effectively lowered by a multi-ion direct knock-on permeation mechanism, which promotes ion occupancy at S4. Replacing S624 with threonine removes this conformational penalty, enhancing outward conduction but prolonging ion retention during inward conduction. This indicates that voltage-dependent interplay between S4 flexibility and multi-ion coordination underlies the characteristic asymmetric conductance in hERG.

### 3.2 hERG Inhibitors Occlude K^+^ Access to the Selectivity Filter

Next, we performed MD simulations of hERG-N629D bound to three classical inhibitors (astemizole, E-4031, and pimozide), using binding poses derived from recent cryo-EM structures (PDB: 8ZYO, 8ZYP, and 8ZYQ).^17^ All three inhibitors occupy a common binding site within the central cavity, between S624 and Y652, below the S4 site of the SF (**Figure 2a**). Astemizole has been reported to adopt two binding modes within the central cavity, differing in hydrogen-bonding interactions at S624 and the orientation of the benzimidazole moiety relative to Y652, with no significant difference in binding free energy between them.^17^ Both binding modes were therefore simulated.

**Figure 2.**
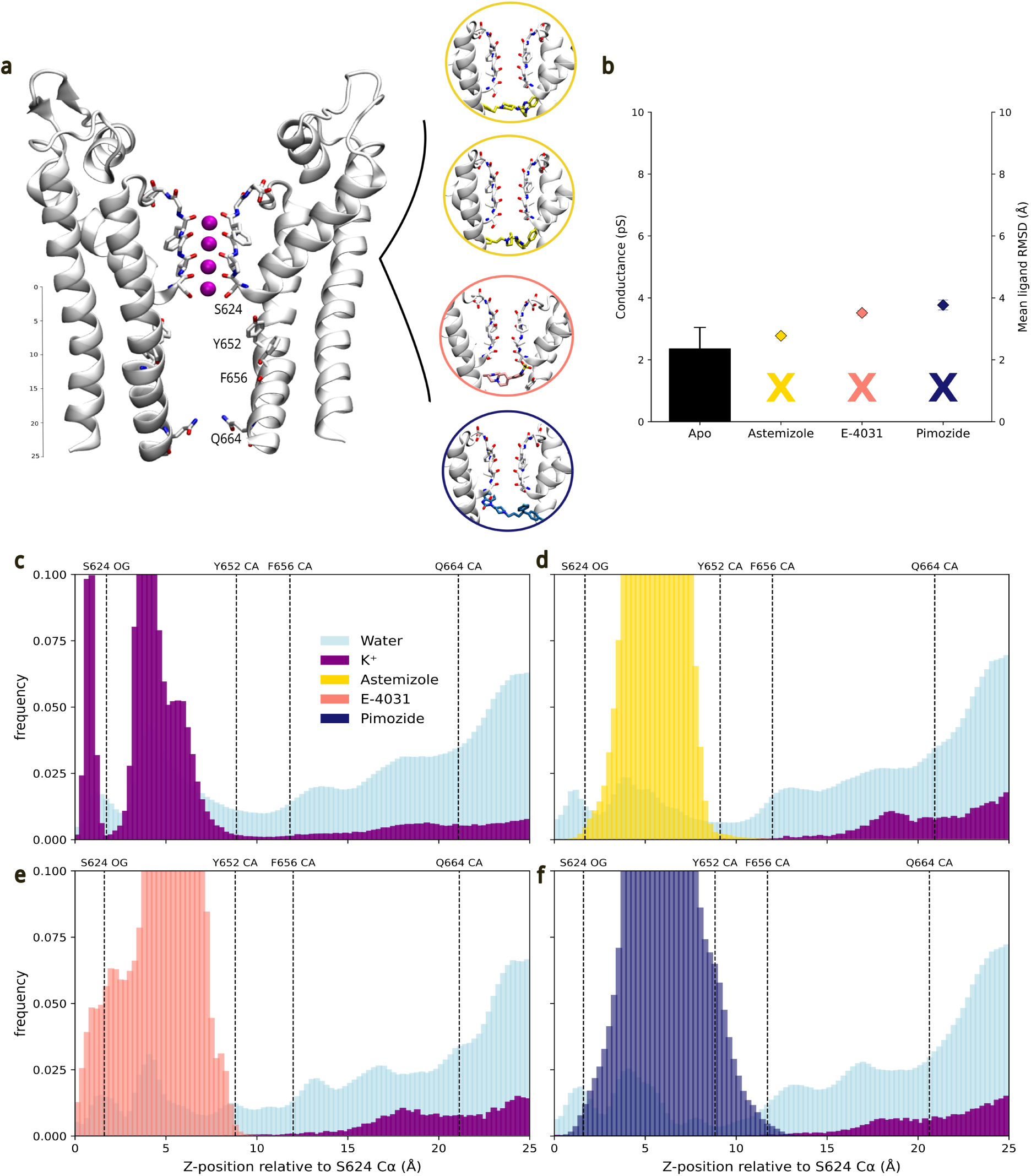
Effect of hERG inhibitors on hERG permeation and cavity occupancy. (a) Representative snapshot of the apo hERG-N629D channel, with key residues S624, Y652, F656, and Q664 labelled. K^+^ ions are shown as purple spheres. Insets show the cryo-EM-derived binding poses of astemizole poses 1 and 2 (yellow), E-4031 (pink), and pimozide (blue) within the central cavity. (b) Single-channel conductance at +300 mV for apo and inhibitor-bound simulations. No K^+^ permeation events were observed in any inhibitor-bound replica, indicated by crosses. Mean ligand RMSD values relative to the cryo-EM pose are shown on the right y-axis as coloured diamonds, confirming stable binding throughout. (c–f) K^+^ (purple) and water (light blue) density distributions along the pore axis relative to the S624 C*α* position for (c) apo, (d) astemizole, (e) E-4031, and (f) pimozide simulations at +300 mV. Dashed vertical lines indicate the positions of key cavity-lining residues.

No K^+^ permeation events were observed in any inhibitor-bound simulation, in contrast to the robust conductance observed in apo simulations under identical conditions (**Figure 2b**). This is consistent with the well-established finding that these inhibitors impede ion movement through the channel’s open state.^30,31^ To verify that the absence of permeation is attributable to the bound inhibitor, we confirmed that all three inhibitors remained stably bound throughout the simulations, with ligand RMSD values remaining low relative to the cryo-EM pose (**Figure 2b**), indicating no ligand displacement over the course of the simulations, in line with their nanomolar binding affinities to hERG.^32^

Structural studies of inhibitor-bound hERG have proposed a “plug-like” occlusion model, in which these compounds suppress K^+^ conduction by occluding the intracellular entrance of the SF, based on inhibitor positions in cryo-EM structures.^17,32^ To directly assess the consequence of this binding on ion permeation, we measured ion and water density distributions along the pore axis. In apo simulations, a continuous ion density is observed along the pore domain with a pronounced peak at SCav (*z ∼* 2–9 Å) (**Figure 2c**). As all three compounds occupy this region, this peak is abolished in all inhibitor-bound systems (**Figure 2d-f**), supporting the plug-like occlusion mechanism whereby bound inhibitors physically block ion access to the SF. Water density within the central cavity, however, remains comparable to that observed in apo simulations, indicating that inhibitor binding does not cause additional dehydration of the cavity (**Figure 2c-f**). Taken together, our simulations capture the dynamic effect of inhibitor binding, reinforcing the view that these compounds act as molecular plugs that sterically occlude K^+^ access to the SF.

To further characterise inhibitor binding modes, we used ProLIF^33^ to quantify protein-ligand interactions across all inhibitor-bound systems (**Figure S2a-d**). Consistent contacts were identified with L622, T623, S624, G648, Y652, and F656, in good agreement with both mutagenesis^34–37^ and structural data.^17^ Y652 showed the highest interaction frequencies across all compounds, with *π*–*π* stacking and cation–*π* interactions observed consistently across all four subunits (**Figure S2a-d**). Y652 is a primary determinant of inhibitor binding in hERG.^35,38^ Notably, the role of F656 has been debated, with early studies supporting a direct role,^36,39^ but our analysis shows that contacts were present with lower and less consistent frequencies relative to Y652, in better alignment with the recent proposal that F656 may contribute more allosterically rather than through direct contact.^17^

Finally, we examined whether inhibitor binding induced any structural perturbations to the SF. Previous structural studies compared static cryo-EM structures of apo and inhibitor-bound hERG, finding no significant conformational changes upon inhibitor binding.^32^ Whether this holds under applied voltage over extended simulation timescales remains unresolved. We find no significant structural perturbation of the SF across all inhibitor-bound systems (**Figure S3a-e**). Two K^+^ ions occupied sites S2 and S3 of the SF at the start of all inhibitor-bound simulations and remained stably bound throughout, confirming that inhibitor binding does not perturb SF ion occupancy. The suppression of ion permeation is therefore a consequence of physical occlusion by the bound inhibitor, rather than any inhibitor-induced conformational change of the SF such as carbonyl rotation, which has been demonstrated to impede ion permeation in hERG.^27,40^ Together, this indicates that hERG inhibition operates through physical occlusion of the permeation pathway, rather than triggering a transition to non-conductive SF geometries.

### 3.3 Fenestration-bound activators enhance K^+^ permeation by increasing cavity ion occupancy

Negatively charged activators (NCAs) increase channel currents across a range of K^+^ channels, including K2P and BK channel families.^19^ Whether hERG NCAs such as PD-118057 and PD-307243 activate hERG via the same conductance-enhancing mechanism or instead act by attenuating inactivation and/or shifting voltage dependence remains a topic of debate. One proposed site positions these NCAs within the central cavity beneath the SF (**Figure S4a**), where the negatively charged group has been proposed to coordinate a K^+^ ion to promote channel opening,^19^ while an alternative mode places the activator within a side fenestration pocket of the cavity, consistent with mutagenesis data identifying F619 and L646 as key determinants of activator binding (**Figure 3a**).^20,41^ We used our hERG-N629D system to investigate these two competing binding modes and assess their effects on ion permeation.

**Figure 3.**
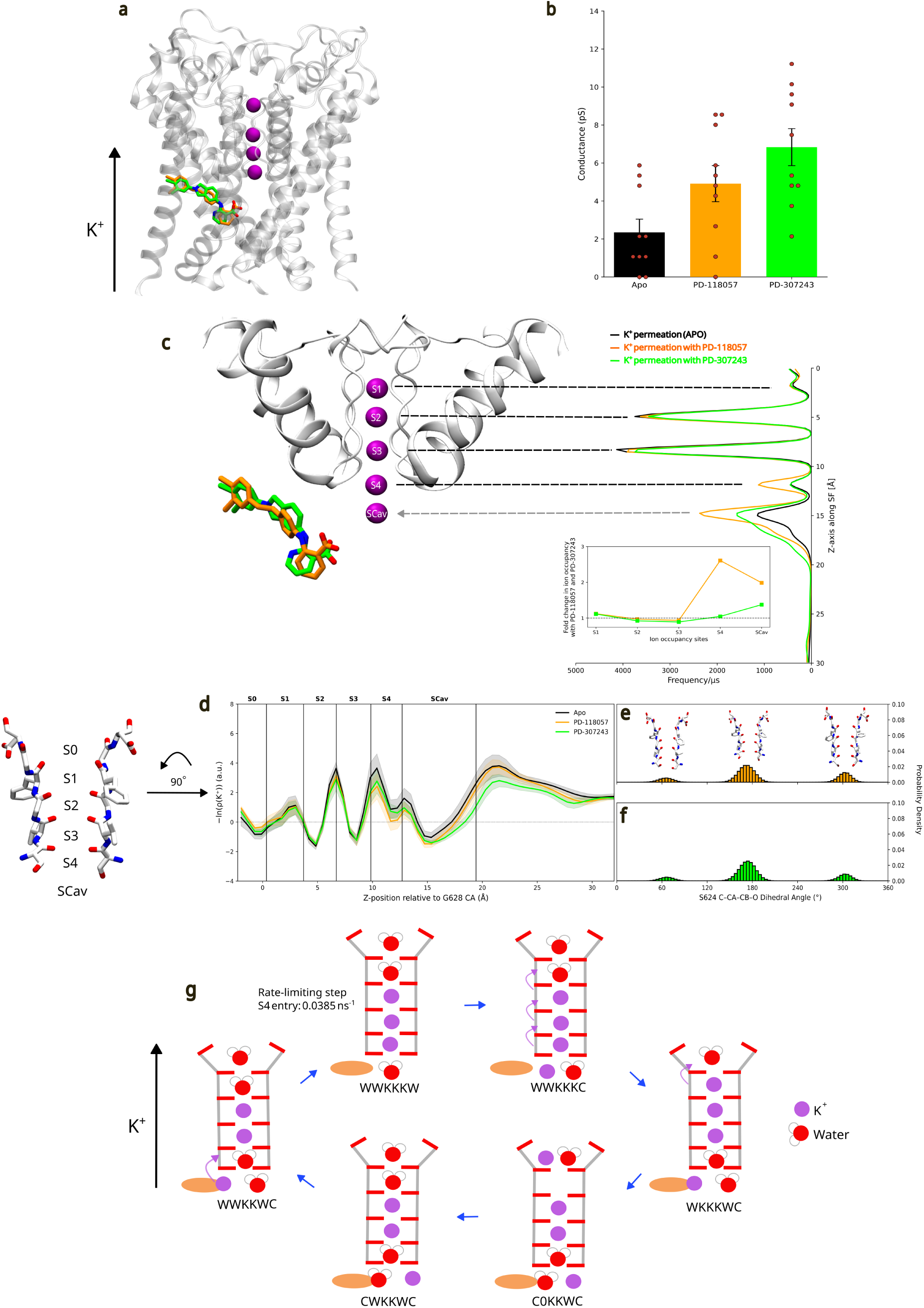
Effects of fenestration-bound PD-118057 and PD-307243 on pore K^+^ occupancy and permeation. (a) Snapshot of hERG-N629D with PD-118057 (orange) and PD-307243 (green) overlaid at the fenestration site of the central cavity. K^+^ ions are shown as purple spheres. (b) Single-channel conductance values for apo, PD-118057, and PD-307243 simulations at +300 mV. (c) Ion occupancy (frequency/*µ*s) of K^+^ binding sites (S0 to SCav) obtained from permeation MD simulations of hERG-N629D in the presence or absence of PD-118057 and PD-307243. The grey arrow points to the increase in SCav K^+^ occupancy. The inset shows the fold change in ion occupancy across S1 to SCav sites. (d) PMF profiles of K^+^ along the pore axis for apo (black), PD-118057 (orange), and PD-307243 (green), with shaded regions indicating standard error. SF and SCav binding site positions are indicated by vertical lines. (e,f) Probability density distributions of the S624 C-C*α*-C*β*-O dihedral angle for PD-118057 (e, orange) and PD-307243 (f, green), with representative rotameric conformations shown above each distribution. (g) Dominant permeation cycle of hERG-N629D in the presence of PD-118057 at +300 mV resolved by Kperm analysis. Each state represents the occupancy of binding sites S0–S4 and SCav, where K denotes K^+^, W denotes water, C denotes co-occupancy, and 0 denotes an empty site. K^+^ ions are shown as purple spheres, water as red spheres, and the fenestration-bound activator as orange ellipses. The rate-limiting step is ion entry into S4 from SCav, with a rate of 0.0385 ns*^−^*^1^. Error bars represent SEM.

When placed within the central cavity, little to no K^+^ permeation was observed for either compound (**Figure S4b**). Instead, their behaviour was more reminiscent of inhibitor binding; therefore, it is difficult to reconcile this binding mode with channel activation. These findings indicate that the central cavity is unlikely to represent the functionally relevant binding site for either compound. In contrast, fenestration-bound PD-118057 and PD-307243 induced pronounced effects on hERG ion permeation. First, channel conductance increased relative to the apo channel, reaching 4.91 *±* 0.95 pS for PD-118057 and 6.84*±*0.97 pS for PD-307243 (*∼*2.1-fold and *∼*2.9-fold increases, respectively; Welch’s *t*-test: apo vs PD-118057, *p* = 0.044; apo vs PD-307243, *p* = 0.0017) (**Figure 3b**). Second, K^+^ occupancy at the SCav site increased by approximately two-fold for PD-118057 and *∼*1.4-fold for PD-307243 (**Figure 3c**). The effect on S4 occupancy differed slightly between the compounds, with PD-118057 inducing a *∼*2.5-fold increase whereas PD-307243 showed little change relative to apo (**Figure 3c**). We hypothesise that this may be explained by the higher permeation rate observed with PD-307243, where more rapid K^+^ transit through S4 would result in lower time-averaged occupancy at the S4 site. Collectively, these results indicate that fenestration-bound hERG activators enhance K^+^ permeation by increasing ion occupancy below the SF.

These observations are consistent with those reported for a related NCA, BL-1249, which has been shown to increase macroscopic currents in hERG and unitary conductance in K2P channels.^19^ MD simulations of BL-1249 in TRAAK revealed a *∼*16-fold increase in K^+^ occupancy below the SF and increases at the S1 and S4 binding sites, accompanied by a 1.6-fold increase in K^+^ permeation rate, a mechanism driven by stabilisation of K^+^ ions below the SF by the NCA.^19^ Our simulations reveal a similar pattern, with increased K^+^ occupancy at the SCav site for both fenestration-bound PD-118057 and PD-307243, alongside enhanced outward K^+^ permeation. Single-channel conductance measurements have been reported for PD-118057 but not for PD-307243. To the best of our knowledge, the experimental measurements have only been performed at negative voltages, and while the current magnitude largely agrees, PD-118057 does not increase single-channel conductance.^41^

Finally, we examined the effects of activator binding on the free energy landscape and permeation kinetics. The increased outward conductance is accompanied by a reduction in the free energy barriers at the SCav-to-S4 and S4-to-S3 transitions relative to apo (**Figure 3d**), supporting the idea that increased ion availability below the SF facilitates outward permeation. Given that stabilisation of S624 into a more conductive rotameric state underlies the increased outward permeation observed in the S624T mutant, we examined whether activator binding similarly influences S624 dynamics. Dihedral analysis reveals that the rotameric distribution of S624 in the presence of fenestration-bound activators closely resembles that of the apo channel (**Figure 3e,f**), indicating that the increase in conductance is not attributable to direct stabilisation of S624 into a more conductive geometry. Permeation cycle analysis of PD-118057 reveals that the rate-limiting step remains ion entry into S4 from SCav, but this transition proceeds approximately 2.5-fold faster (**Figure 3g**). We attribute this acceleration to increased K^+^ availability at SCav: with more ions populating the cavity, productive entry attempts into S4 occur more frequently, thereby enhancing outward conductance. While the functional outcome is consistent with that reported for other NCAs, we next investigated whether these binding poses are supported by experimental mutagenesis data.

### 3.4 Fenestration-bound activators adopt stable binding poses consistent with mutagenesis data

To characterise ligand dynamics and binding-site interactions in greater detail, we performed protein-ligand interaction fingerprinting and ligand RMSD analysis across all production replicas for both proposed binding sites. Experimentally, scanning mutagenesis of PD-118057 identified F619 and L646 as the primary structural determinants of agonist activity, with the compound proposed to occupy a hydrophobic site between two adjacent subunits formed by these residues.^41,42^ Computational docking of PD-307243 similarly supports fenestration site binding, with contacts at F619 and L646.^20^ F557 has additionally been implicated in activator binding.^42^ Direct contacts with these residues were observed in the starting structures of both compounds at the fenestration site (**Figure 4a-c**). Protein-ligand interaction fingerprinting confirmed that F619, L646, and F557 were among the residues with the highest interaction frequencies for both compounds across all replicas (**Figure 4d,e**). In contrast, when the activators were bound in the central cavity, these interactions were absent, and instead the activators were in contact with Y652 and F656 (**Figure S5a-c**). These interactions remained dominant through-out the production replicas (**Figure S5d,e**), which is inconsistent with mutagenesis data demonstrating that Y652 has minimal effect on activator binding.^39,43^

**Figure 4.**
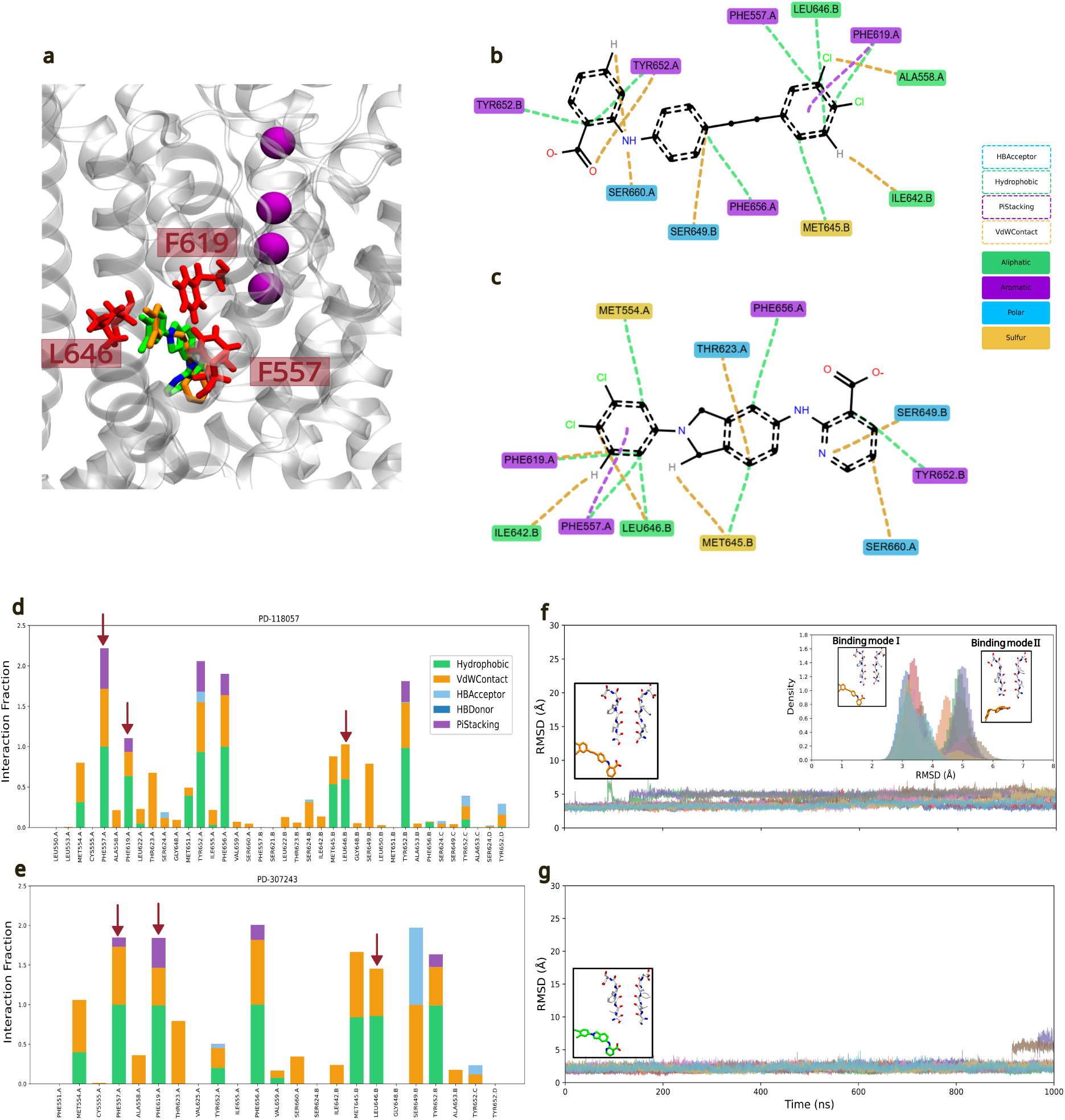
Protein-ligand interactions and binding stability of fenestration-site activators. (a) Representative snapshot of the hERG-N629D fenestration site showing key interacting residues F557, F619, and L646 in red licorice representation, with PD-118057 (orange) and PD-307243 (green) overlaid. K^+^ ions are shown as purple spheres. (b,c) 2D interaction diagrams for PD-118057 (b) and PD-307243 (c). (d,e) Protein-ligand interaction fraction for PD-118057 (d) and PD-307243 (e), averaged across all MD simulations at +300 mV. Red arrows highlight interactions with F557, F619, and L646. (f) Per-replica RMSD of PD-118057 over 1 *µ*s MD simulations at +300 mV, wrt initial minimised structure. Bottom-left inset: initial minimised structure of PD-118057. Top-right inset: RMSD distributions across all replicas, with representative structures for binding mode I and binding mode II. (g) Per-replica RMSD of PD-307243 over 1 *µ*s MD simulations at +300 mV, wrt the initial minimised structure. Bottom-left inset: initial minimised structure of PD-307243.

We assessed the stability of both binding poses by computing ligand RMSD across all production replicas. Fenestration-bound PD-118057 and PD-307243 remained stably bound throughout, with RMSD values consistently below 6 Å and no unbinding observed across any replica (**Figure 4f,g**). In contrast, central cavity binding was markedly less stable, with two of five PD-118057 replicas and one of five PD-307243 replicas showing RMSD values exceeding 10 Å, indicative of ligand unbinding (**Figure S5f,g**). The fenestration-site stability observed here is consistent with the findings for another hERG activator, ICA-105574, which remained stably bound at the same inter-subunit fenestration site over 500 ns of MD simulations.^44^ Notably, PD-118057 adopted two distinct binding geometries within the fenestration (**Figure 4f**): three of ten fenestration-bound replicas retained the initial starting pose (Binding mode I), while the remaining replicas converged to an alternative orientation (Binding mode II), characterised by a kinked hydrophobic tail and reorientation of the carboxylate-bearing aromatic ring. Despite these structural differences, both modes remained stable throughout. PD-307243, in contrast, adopted a single binding geometry across all replicas (**Figure S6**).

## 4. Conclusion

In this work, we combined the N629D SF-stabilising mutant with ECC-based force fields to enable robust *in silico* electrophysiology simulations of hERG. Given the flexible, non-canonical SF of hERG, which has previously prevented sustained ion permeation, our approach not only reproduces experimental single-channel conductance and its asymmetric characteristics for the first time, but also unlocks quantitative, physics-based simulations of hERG.

Our findings can be summarised in three key points. First, the combination of ECC force fields and the N629D mutation resolves the longstanding challenge of sustaining K^+^ permeation in hERG simulations, with conductance values and voltage-dependent asymmetry in agreement with experiment, and permeation proceeding via a direct knock-on mechanism without co-permeating water molecules. Second, the conformational flexibility of S624, which lines the S4 ion-binding site, controls asymmetric conductance in hERG. S624 frequently samples non-conductive rotameric geometries, whereas S624T eliminates conductance asymmetry in our simulations, strongly indicating that S624 dynamics represent a structural determinant of this single-channel conductance asymmetry. Third, simulations of hERG inhibitors and activators provide mechanistic insight into small-molecule modulation. Inhibitors abolish permeation through direct steric occlusion of K^+^ access to the SF, stabilised through direct contacts with hydrophobic residues lining the central cavity. In contrast, fenestration-bound activators enhance K^+^ permeation and SF ion occupancy, with PD-118057 and PD-307243 producing two- to threefold increases in conductance. These results support the previously proposed fenestration binding mode and are consistent with mutagenesis data (**Figure 5**).

**Figure 5.**
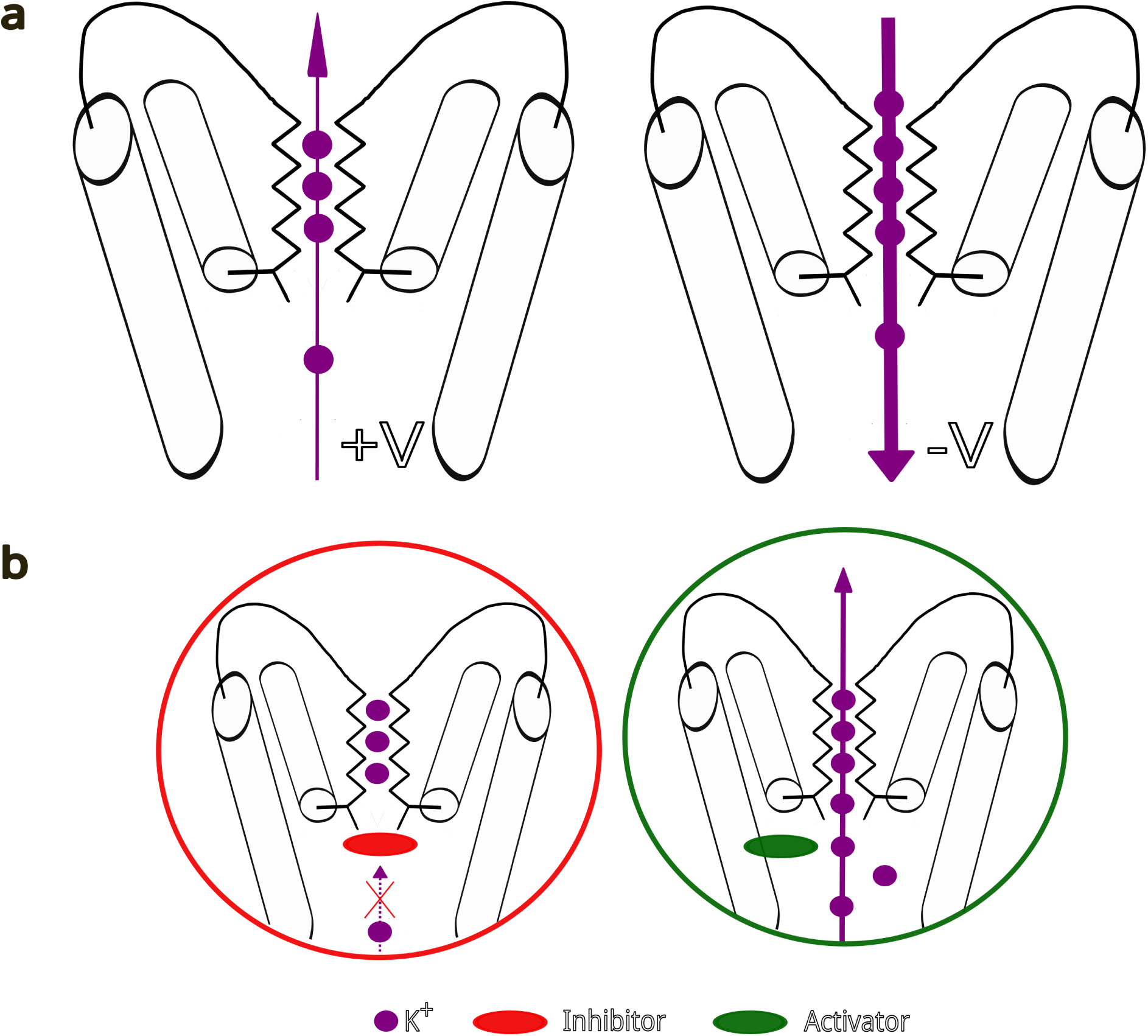
Summary of asymmetric conductance and proposed mechanisms of cavity inhibition/activation. (a) S624 site frequently samples non-conductive rotameric geometries, limiting S4 occupancy and outward conductance at positive voltage (+V). At negative voltage, incoming K^+^ ions are driven through S4 regardless of its geometry, resulting in the characteristic asymmetry in hERG single-channel conductance. (b) Inhibitors (red) bound in the central cavity directly block K^+^ access to the SF, abolishing permeation. In contrast, activators (green) bind in the fenestration site, increasing K^+^ occupancy and enhancing outward K^+^ permeation. Arrow width indicates the relative magnitude of K^+^ permeation.

The simulation framework established here provides a basis for assessing hERG modulation beyond binding affinity or potency alone. While these measures describe how strongly a compound interacts with hERG, they do not directly describe the functional consequences of binding. By enabling the effects of small molecules on ion permeation to be resolved directly, our approach provides an additional functional readout of hERG modulation at the molecular level.

## 5. Methods

### 5.1 System Preparation and Protocol

#### 5.1.1 hERG-N629D apo

The atomistic model of the hERG channel was based on the Protein Data Bank (PDB) entry 9CHP.^40^ Four K^+^ ions, obtained from the MthK crystal structure,^45^ were positioned within the S1–S4 binding sites of the selectivity filter. Using CHARMM-GUI, the protein was embedded in a POPC membrane using the OPM database and solvated with aqueous KCl.^46^ The initial system consisted of one homotetrameric channel, 535 POPC molecules, 68407 water molecules, 636 K^+^ ions, and 636 Cl*^−^* ions, corresponding to a 0.5 M salt concentration.

To generate the mutant system, residue N629 was aligned with D447 of the Shaker channel (PDB ID: 7SIP),^47^ and the asparagine side chain was manually replaced with the aspartate side chain. The final system consisted of one channel, 535 POPC molecules, 68411 water molecules, 636 K^+^ ions, and 632 Cl*^−^* ions, corresponding to a 0.5 M salt concentration. Ten independent 1 µs simulations were performed at each voltage (+300 and *−*300 mV).

#### 5.1.2 hERG-N629D with S624T mutation

The hERG-N629D system was used as the starting model. Residue S624 was aligned with T442 of the Shaker channel (PDB ID: 7SIP),^47^ and serine residues were manually replaced with threonine residues. The final system consisted of one channel, 535 POPC molecules, 68411 water molecules, 636 K^+^ ions, and 632 Cl*^−^* ions, corresponding to a 0.5 M salt concentration. Ten independent 1 µs simulations were performed at each voltage (+300 and *−*300 mV).

#### 5.1.3 hERG-N629D with inhibitors bound

The hERG-N629D system was used as the starting model. A representative frame from our apo simulation was selected based on the lowest mean RMSD of central cavity residues across all three inhibitor-bound cryo-EM structures (PDB ID: 8ZYO, 8ZYP, and 8ZYQ).^17^ The selectivity filter of the cryo-EM structures was aligned to this selected frame, and ligand coordinates were extracted to generate the initial binding poses. Hydrogen atoms were added to each ligand, and protonation states were assigned at pH 7.4 using Open Babel.^48^

Four inhibitor-bound systems were constructed: astemizole (pose 1 and pose 2, based on distinct binding orientations reported in the literature), E-4031, and pimozide. Each system consisted of one tetrameric channel, 535 POPC molecules, 68410 water molecules, 636 K^+^ ions, and 633 Cl*^−^* ions, corresponding to a 0.5 M salt concentration. Five independent 500 ns replicas were run for each ligand pose at +300 mV.

#### 5.1.4 hERG-N629D with activators bound (central cavity)

Using the same representative frame selected for the inhibitor-bound systems, the receptor was prepared for docking using the Dock-Prep pipeline, which included the addition of polar hydrogens, assignment of Gasteiger charges, and conversion to PDBQT format.^49^ PD-118057 and PD-307243 were prepared from SMILES representations with the carboxylate group in the deprotonated state. Polar hydrogens were added, and 3D coordinates were generated using RDKit;^50^ the resulting structures were converted to PDBQT format using Meeko.^51^ Docking was performed using AutoDock Vina,^52,53^ with a box size of 20 *×* 20 *×* 20 ^Å^, covering the central cavity below the selectivity filter. An exhaustiveness of 32 was used, generating 20 poses per docking run across three independent runs, yielding 60 poses per compound in total. The selected pose was converted to PDB format, and hydrogen atoms were added with protonation states assigned at pH 7.4 using Open Babel.^48^ The pose showing the closest agreement with activator binding modes reported in the literature was selected as the initial structure for MD simulations. Each system consisted of one tetrameric channel, 535 POPC molecules, 68410 water molecules, 637 K^+^ ions, and 632 Cl*^−^* ions, corresponding to a 0.5 M salt concentration. For the cavity binding mode, five independent 1 *µ*s simulations per compound were carried out at +300 mV.

#### 5.1.5 hERG-N629D with activators bound (fenestration site)

The protein conformation used for docking was taken from an unrestrained MD simulation in which the fenestration site was observed to be open. The protein and ligands were prepared as described in Section 5.1.4. Docking was performed using AutoDock Vina^52,53^ with a box size of 20 *×* 20 *×* 20 ^Å^ centred at coordinates (92.35, 75.51, 73.07) ^Å^, which covers the fenestration site adjacent to the cavity. An exhaustiveness of 32 was used, generating 20 poses per docking run across three independent runs, yielding 60 poses per compound in total. Following docking, the selected pose was converted to PDB format, and all hydrogen atoms were added with protonation states assigned at pH 7.4 using Open Babel.^48^ The pose showing the closest agreement with fenestration site binding modes reported in the literature was selected as the initial structure for MD simulations.^20^ Each system consisted of one homotetrameric channel, 535 POPC molecules, 68410 water molecules, 637 K^+^ ions, and 632 Cl*^−^* ions, corresponding to a 0.5 M salt concentration. For the fenestration binding mode, ten independent 1 *µ*s simulations were performed for each compound at +300 mV.

### 5.2 Force field-specific parameters

We applied a recently developed MD force field optimised for K^+^ channels in our simulations.^24^ Partial charges of ionised groups, including the side chains of arginine, lysine, glutamate and aspartate, as well as free ions, were rescaled to yield charges of *±*0.78*e*.

#### 5.2.1 Simulation parameters for Amber14SB-rescaled

The Amber14SB FF was sourced from the GROMACS user contributions repository. For the Amber14SB force field, the Lipid21^54^ and TIP3P^55^ water models were used. Key topology files were modified, including a reduction of the carbonyl oxygen charge at the S3 site from *−*0.55670 to *−*0.53, with the remaining charge redistributed across the given residues—a correction shown to enhance conductance.^24^ van der Waals (vdW) interactions used a 1.0 nm cut-off with analytical dispersion corrections, and long-range electrostatics were treated with PME with a 1.2 nm cut-off.

#### 5.2.2 Simulation parameters for ligands

Ligand parametrisation was handled using the pmx Python package.^56^ Ligands were parametrised with GAFF 2 using ACPYPE^57^ and Antechamber^58^ with AM1-BCC^59^ charges and off-site charges for halogen atoms.^60^ In line with the Electronic Continuum Correction (ECC) approach,^24^ ligand atomic charges were redistributed and scaled to yield a net charge of +0.78*e* for inhibitors and *−*0.78*e* for activators, accounting for the lack of explicit electronic polarisation in classical force fields.

### 5.3 Molecular Dynamics (MD) simulations

All simulations were performed with GROMACS, versions 2022.2–2025.1.^61^ The starting structure was minimised using the steepest descent algorithm until the maximum force was below 1000 kJ mol*^−^*^1^ nm*^−^*^1^. Mild distance restraints (1000 kJ mol*^−^*^1^ nm*^−^*^2^) were applied between Y616 and D629 to prevent collapse of the SF, which has been shown to be an important determinant of filter stability in hERG and is analogous to conserved hydrogen bond networks in other K^+^ channels.^27,62^ Distance restraints between cross-facing F656 and R665 C*α* atoms were also applied to avoid narrowing of the lower gate, which would otherwise restrict ion and water entry into the central cavity (**Figure S7a– d**).^63^

A 500 ps NVT equilibration was performed with harmonic restraints of 1000 kJ mol*^−^*^1^ nm*^−^*^2^ applied to all heavy atoms of the protein and POPC lipids, and ligands where applicable. Temperature was maintained using the v-rescale thermostat.^64^ This was followed by a 1750 ps NPT equilibration using the same restraint strength, with pressure semiisotropically coupled to 1 bar using the c-rescale barostat.^65^ Subsequently, a 100 ns NPT equilibration was conducted during which all position restraints were removed. A time step of 2 fs was used, and constraints were applied to bonds involving hydrogen using the LINCS algorithm.^66^

To study ion permeation, we used an applied electric field along the *z*-axis across a POPC bilayer membrane. The box *z* dimension in the final frame of the 100 ns equilibration was used to set the electric field. Transmembrane voltage, *V*, was calculated using:

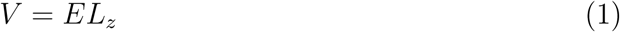

where *E* is the applied electric field in V/nm and *L_z_* is the length of the box along the *z*-axis in nm.^67^ Production runs were conducted at applied voltages of +300 or -300 mV, which were found earlier to yield a good balance between simulation efficiency and minimal perturbation of the channel.^68^ Simulation lengths for each system are specified in Sections 5.1.1–5.1.5.

## 6. Author Contributions

V. Gapsys and W. Kopec conceived the project. I. Manoza performed and analysed the MD simulations. I. Manoza, V. Gapsys, and W. Kopec wrote the manuscript.

## 7. Acknowledgements

This work was supported by the UK Research and Innovation (UKRI) Biotechnology and Biological Sciences Research Council (BBSRC) AI for Drug Discovery Programme.

This project made use of time on HPC granted via the UK High-End Computing Consortium for Biomolecular Simulation (HECBioSim; https://www.hecbiosim.uk), supported by the Engineering and Physical Sciences Research Council (EPSRC) (grant no. EP/X035603/1).

