## Supplemental Material for "Computational assay of hERG ion permeation and its modulation by small-molecule compounds"

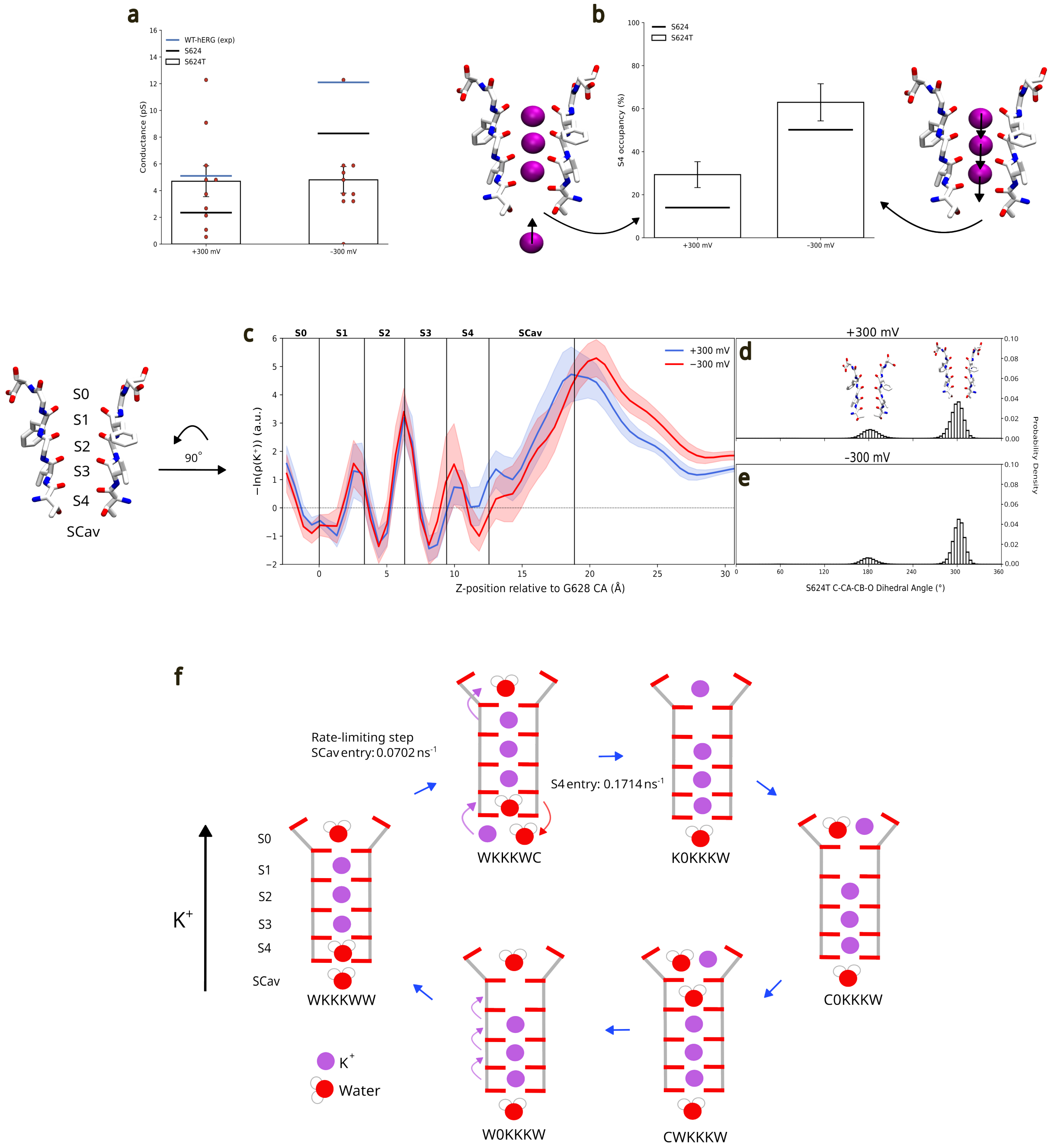
a
WT-hERG (exp)
S624
S624T
Conductance (pS)
+300 mV
-300 mV
b
S624
S624T
S4 occupancy (%)
+300 mV
-300 mV
S0
S1
S2
S3
S4
SCav
90°
c
S0
S1
S2
S3
S4
SCav
+300 mV
−300 mV
−ln(ρ(K+)) (a.u.)
Z-position relative to G628 CA (Å)
+300 mV
d
-300 mV
e
Probability Density
S624T C-CA-CB-O Dihedral Angle (°)
f
Rate-limiting step
SCav entry: 0.0702 ns-1
S4 entry: 0.1714 ns-1
WKKKWC
K0KKKW
C0KKKW
CWKKKW
W0KKKW
WKKKWW
K+
K+
Water

**Figure S1. Effect of S624T on ion permeation.** (a) Single-channel conductance values for S624T (white bars) at +300 mV and −300 mV, with individual replica values overlaid as red dots. The blue dashed line indicates the experimental WT hERG single-channel conductance, and the black line represents the mean conductance of simulated hERG-N629D with WT S624 for reference. (b) S4 ion occupancy for S624T (white bars) at +300 mV and −300 mV. The black line represents S4 ion occupancy of hERG-N629D with WT S4 for reference. Insets show structural cartoons of the selectivity filter at each voltage, as in Figure 1d. (c) PMF profiles of $K^+$ along the pore axis for S624T at +300 mV (blue) and −300 mV (red), with shaded regions indicating standard error. Selectivity filter binding site positions are indicated by vertical lines. (d,e) Probability density distributions of the S624T C-C$\alpha$-C$\beta$-O dihedral angle at +300 mV (d) and −300 mV (e), with representative rotameric conformations shown above each distribution. (f) Dominant permeation cycle of S624T at +300 mV resolved by Kperm analysis, with the same state notation as in Figure 1h. The rate-limiting step shifts to ion entry into SCav, with a rate of 0.0702 $ns^{-1}$. The rate of ion entry into S4 is 0.1714 $ns^{-1}$, a ∼11-fold increase relative to WT S624, consistent with the reduced PMF barrier to S4 entry conferred by threonine.

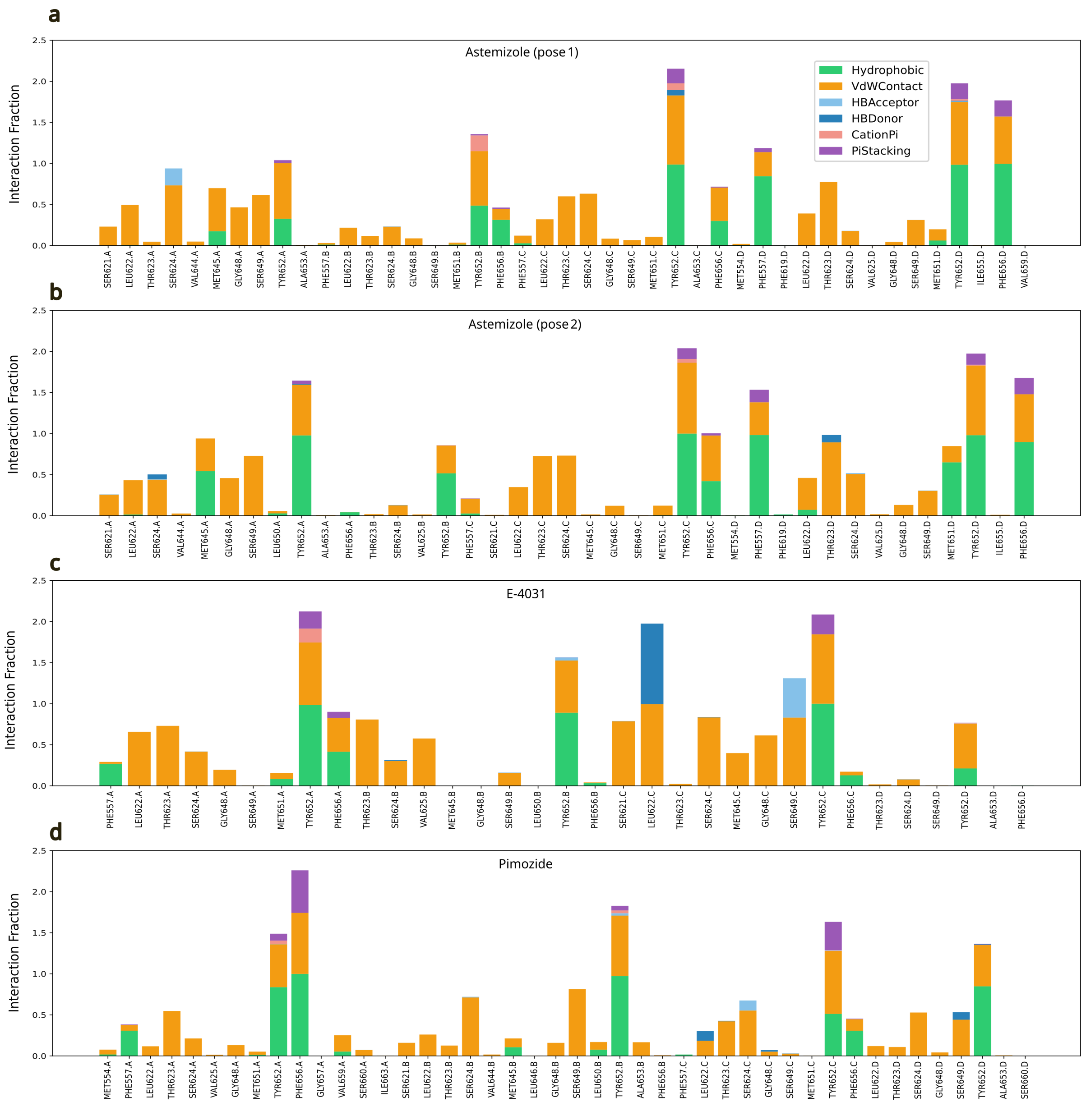


**Figure S2. Protein-ligand interaction fingerprints for inhibitor-bound hERG-N629D simulations.** (a–d) Mean interaction fraction across all production replicas for astemizole pose 1 (a), astemizole pose 2 (b), E-4031 (c), and pimozide (d).

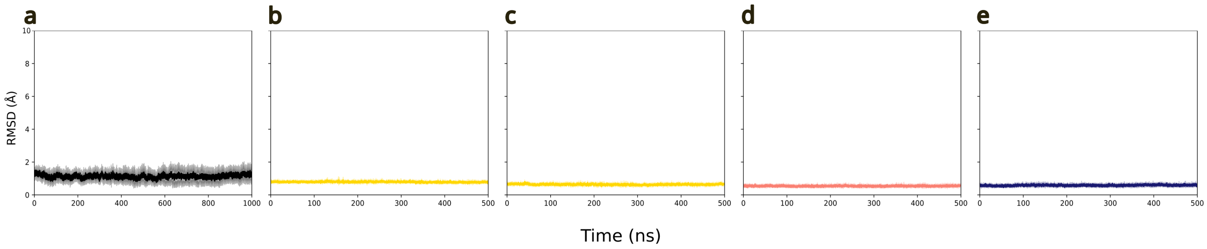


**Figure S3. Selectivity filter stability of apo and inhibitor-bound hERG-N629D simulations.** (a–e) SF backbone RMSD (residues S624–N629) relative to the minimised structure, averaged over all production runs for apo (a, black), astemizole pose 1 (b, yellow), astemizole pose 2 (c, yellow), E-4031 (d, pink), and pimozide (e, blue). Shaded regions indicate standard deviation.

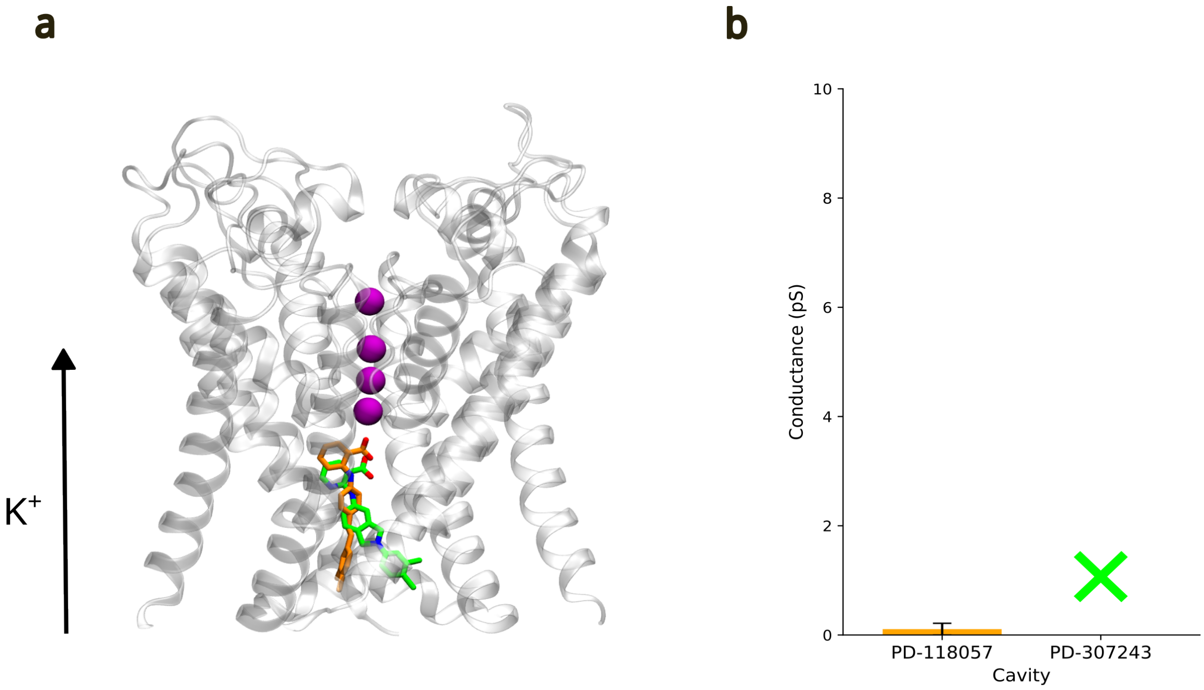


**Figure S4. Effects of central cavity-bound PD-118057 and PD-307243 on $K^+$ permeation.** (a) Snapshot of hERG-N629D with PD-118057 (orange) and PD-307243 (green) overlaid at the central cavity. $K^+$ ions are shown as purple spheres. (b) Single-channel conductance values for PD-118057 and PD-307243 simulations at +300 mV. Error bars represent SEM.

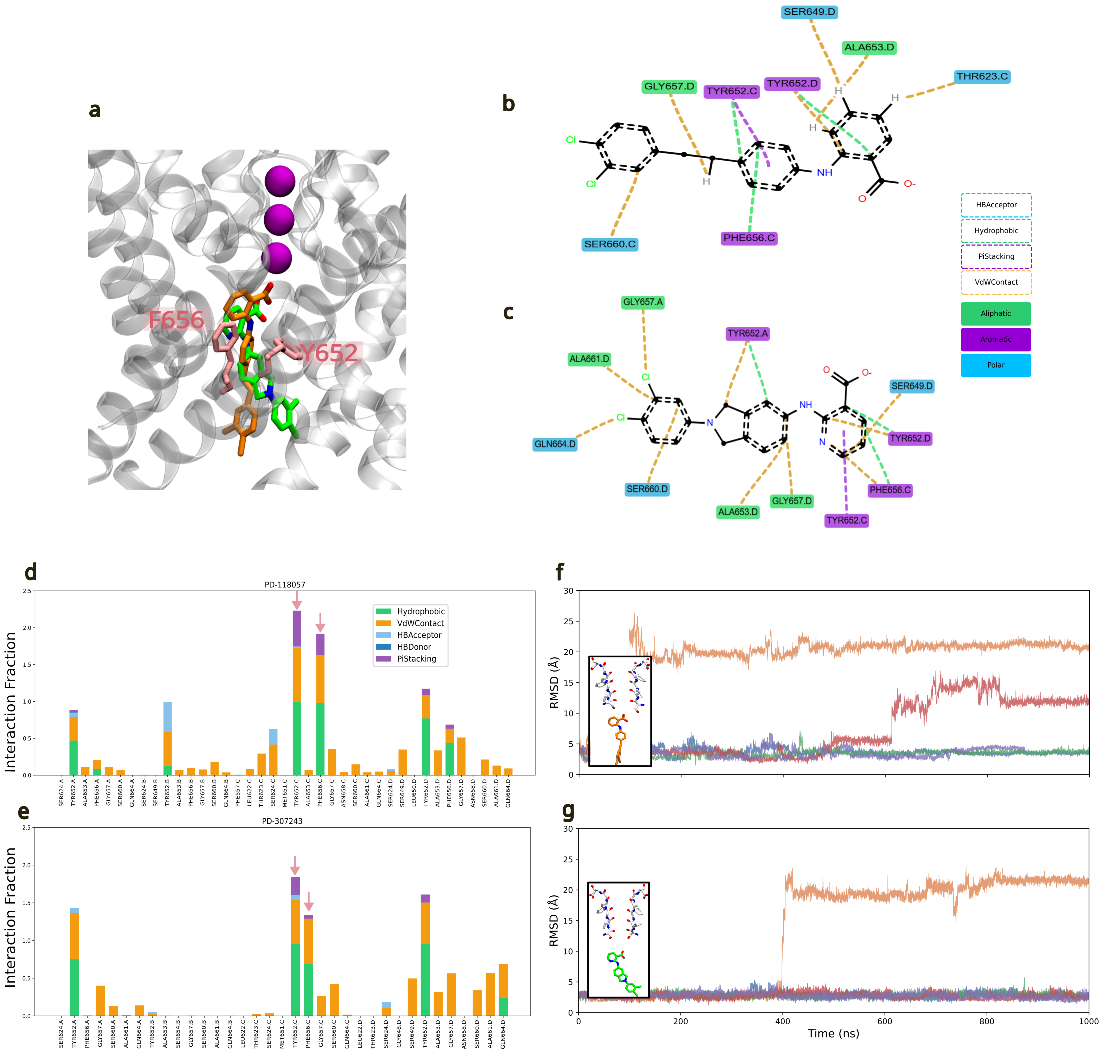

**Figure S5. Protein-ligand interactions and binding stability of central cavity-bound activators.** (a) Representative snapshot of the hERG-N629D central cavity showing key interacting residues Y652 and F656 in pink licorice representation, with PD-118057 (orange) and PD-307243 (green) overlaid. $K^+$ ions are shown as purple spheres. (b,c) Protein-ligand interaction diagrams for PD-118057 (b) and PD-307243 (c). (d,e) Protein-ligand interaction fraction for PD-118057 (d) and PD-307243 (e), averaged across all MD simulations at +300 mV. (f) Per-replica RMSD of PD-118057 over 1 $\mu$s MD simulations at +300 mV, wrt the initial minimised structure. Bottom-left inset: initial minimised structure of PD-118057. (g) Per-replica RMSD of PD-307243 over 1 $\mu$s MD simulations at +300 mV, wrt the initial minimised structure. Bottom-left inset: initial minimised structure of PD-307243.

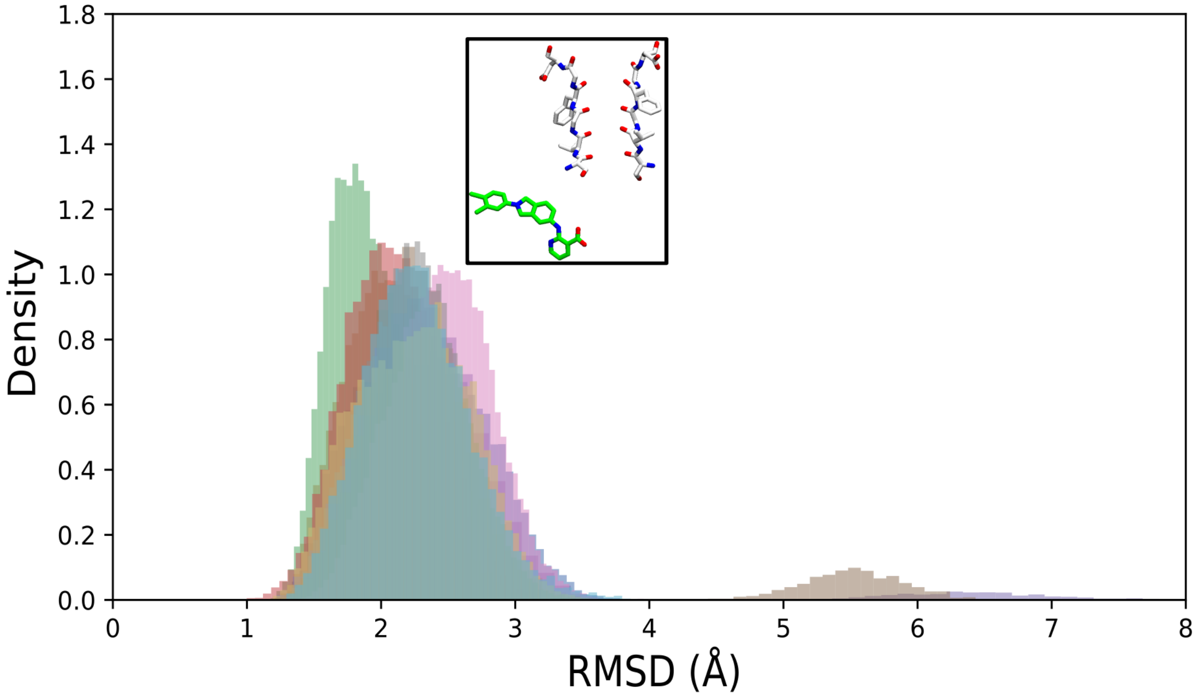


**Figure S6. Ligand RMSD distributions for PD-307243 in the fenestration site.** Each colour represents an individual replica. The inset shows the initial minimised structure of PD-307243.

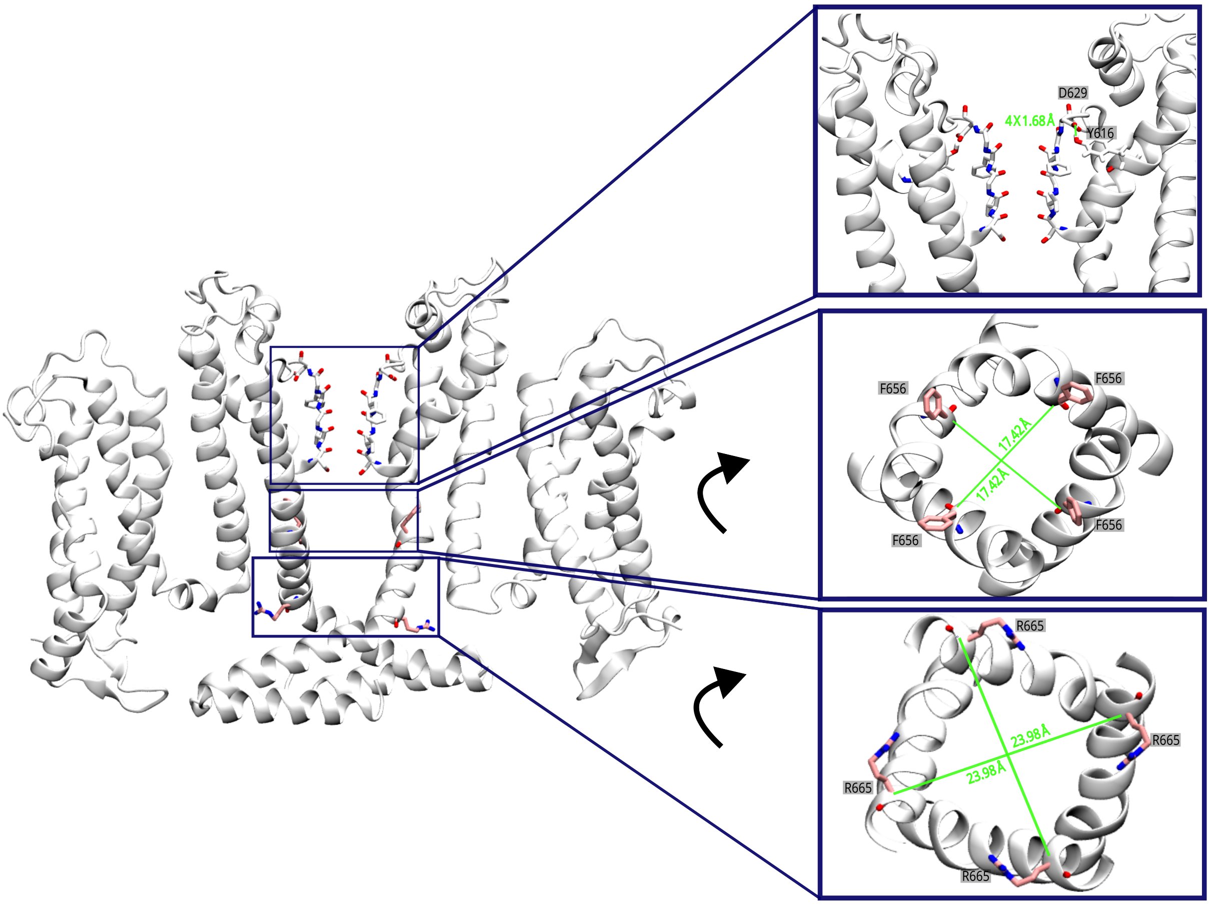


**Figure S7. Distance restraints applied to the hERG channel during MD simulations.** (a) Side view of the transmembrane domain showing two opposite subunits, with the SF, central cavity, and activation gate indicated. (b) Close-up side view of the SF highlighting the distance restraint (green) applied between the O atom of D629 and the hydroxyl hydrogen of Y616, which forms part of the hydrogen-bond network behind the filter. Only one of four equivalent restraints is shown; the restrained distance (green) was taken from the equivalent hydrogen bond between N629 and Y616 in the cryo-EM structure. (c) Top-down view of the central cavity at the level of F656 (pink licorice representation). Distance restraints (green) were applied between the C$\alpha$ atoms of diagonally opposing subunits to maintain the cross-subunit separation observed in the cryo-EM structure. (d) Top-down view of the activation gate at the level of R665 (pink licorice representation). Harmonic distance restraints (green) were applied between the C$\alpha$ atoms of diagonally opposing subunits to maintain the cross-subunit separation observed in the cryo-EM structure.
